# Common variation in *EDN1* regulatory regions highlights the role of PPARγ as a key regulator of Endothelin *in vitro*

**DOI:** 10.1101/2021.11.18.468847

**Authors:** Mauro Lago-Docampo, Carlos Solarat, Luis Méndez-Martínez, Adolfo Baloira, Diana Valverde

## Abstract

Pulmonary Arterial Hypertension (PAH) is a rare disease caused by the obliteration of the pulmonary arterioles, increasing pulmonary vascular resistance and eventually causing right heart failure. Endothelin-1 is a vasoconstrictor peptide whose levels are indicators of disease progression and its pathway is one of the commonest targeted by current treatments.

We sequenced the *EDN1* untranslated regions of a small subset of PAH patients, predicted the effect *in silico*, and used a luciferase assay with the different genotypes to analyze its influence on gene expression. Finally, we used siRNAs against the major transcription factors predicted for these regions (PPARγ, KLF4, and VDR) to assess Endothelin-1 expression in cell culture and validate the binding sites.

First, we detected a SNP in the 5’UTR (rs397751713) and another in the 3’regulatory region (rs2859338) that increased luciferase activity *in vitro* depending on their genotype. We determined *in silico* that KLF4/PPARγ could be binding in the rs397751713 and VDR in rs2859338. By using siRNAs and luciferase, we determined that PPARγ binds differentially in rs397751713. PPARγ and VDR Knock-Down, increased *EDN1* mRNA levels and Endothelin-1 production in PAECs, while PPARγ and KLF4 Knock-Down increased the Endothelin-1 production in HeLa.

In conclusion, common variants in *EDN1* regulatory regions could alter Endothelin-1 levels. We were able to validate that PPARγ binds in rs397751713 and is key to regulate Endothelin-1. Also, KLF4 and VDR regulate Endothelin-1 production in a cell-dependent manner, but for VDR this interaction does not happen by binding directly to the regions we studied.

## Introduction

Pulmonary Arterial Hypertension (PAH) is a rare and devastating disease that involves the thickening of the pulmonary arterioles, this leads to an increase in pulmonary vascular resistance and eventually to right heart failure^1^. The genetic basis of PAH has been slowly uncovered during the past decades, and there are now more than 16 genes involved with different degrees of evidence^2^. However, some genes are not causal but may modulate the development or the evolution of the disease. The *Endothelin-1* gene (*EDN1*) may be one of those. The effects of this peptide are widely implicated in the disease and its treatment, as Endothelin receptor antagonists (ERAs) are used to treat the progression of PAH^3^.

The family of endothelins (ET) is constituted by three isoforms of 21 amino acids, which are well known since the final years of the 20^th^ century^4^. ET-1 is the predominant isoform and the most important by its biological function. It is an endogenous and strong vasopressor synthesized mainly by the vascular endothelium. Most ET-1 is released towards the smooth muscle cells in a paracrine/autocrine way, and pulmonary circulation is its main clearance site^5^. However, it can be detected in plasma or serum and its levels are correlated with the severity of certain diseases such as congestive heart failure and Pulmonary Arterial Hypertension (PAH)^6^.

The disruption of the balance between vasoconstriction and vasodilation triggers a wide variety of pathologies in different organs and tissues. Pulmonary vessels are one of the primary targets of ET-1. Plasma levels of ET-1 have been correlated with the severity of PAH and its prognosis, especially when associated with the plasma levels of ET-3, as the ET-1/ET-3 ratio has proven to be a powerful prognostic indicator ^7–9^. The enhanced activity of the endothelin system has been implicated in PAH severity for a long time^10^, but the amount of experimental data demonstrating it is still limited^11^.

In this work, we looked for variants in the UTR regions of *EDN1* to analyze its influence on gene regulation.

## Material and Methods

### Cohort description

The cohort was composed of 36 patients of group I PH: 21 IPAH and 15 Associated PAH (10 connective tissue disease, 3 HIV, and 2 congenital heart disease). Most of the patients had been described in previous studies from our group^12^.

### Mutational screening

We extracted the genomic DNA from peripheral blood using the FlexiGene DNA Kit (Qiagen) according to the manufacturer’s protocol. DNA amplification was performed with 50 ng of genomic DNA from each individual by Polymerase Chain Reaction (PCR) using the NZYtaq II Green Master Mix (NZYtech). The primers used for each of the EDN-1 gene regions are described in Supp. Table 1. The amplification conditions were 95° C for 2 minutes, 35 cycles of 1 minute at 95° C, 30 seconds at each couple of primers annealing temperature (Supp. Table 1) and 1 minute at 72° C; followed by 5 minutes at 72° C. PCR products were separated by electrophoresis through 1% or 2% agarose gels stained with Ethidium Bromide. To confirm fragment length, Brightmax 500-10 Kb DNA ladder (Canvax) and NZYDNA ladder V (NZYtech) were used as molecular weight markers. PCR fragments were purified using ExoSAP-IT kit (USB Corporation, USA), and sequencing was carried out in the C*entro de Apoio Científico-Técnico á Investigación* (CACTI) of the Universidade de Vigo. Every sample was sequenced independently in both forward and reverse strands to confirm the results obtained. Lastly, sequences were aligned to the reference ENSEMBL DNA sequence [ENST00000379375.5].

### *In silico* effect and conservation analyses

The transcription factors were determined using the online software MatInspector (Genomatix)^13^. This bioinformatics software provides information on transcription factors capable of binding to the genomic regions analyzed. Also, we looked for regulatory motifs and *EDN1* conservation using the UCSC genomebrowser^14^.

### Design of the luciferase constructs

To analyze the 5’ UTR variants we used the pGL3-Basic (Promega), amplification of the 1.4 kb of the 5’ UTR region was carried out using Phusion High Fidelity polymerase (ThermoFisher) with the primers shown in Supp. Table 2. Fragment and plasmid were digested with NheI and XhoI (NZYtech), ligated with T4 ligase (Canvax, Spain), and used to transform NZYstar competent cells (NZYtech). The empty pGL3-Basic was used as a negative control and pRL-CMV (Promega) was used as an internal control.

For the 3’ variants, we used the pmirGLO dual-luciferase vector (Promega). A fragment of 1.4 kb was amplified and cloned as stated before but using SalI instead of XhoI. Empty vector was used as the positive control.

### Cell culture and transfection

HeLa cells (ATCC, USA) were cultured in DMEM (ThermoFisher) supplemented with 10 % Fetal Bovine Serum (FBS) (ThermoFisher), 1% streptomycin/penicillin (Lonza), at 37° C with 5% CO_2_ and humidified atmosphere. PAEC (ECACC, United Kingdom) were cultured in Endothelial Cell Growth medium (Sigma-Aldrich) supplemented with 10 % FBS and 1 % streptomycin/penicillin. PAECs were used in passages between 3 and 7.

For the luciferase assay, we plated 40.000 HeLa cells in 24-well plates, at least 4 replicates per condition were used and analyzed on different days. When the cells showed an 80-90% confluence, transfection was carried out using 0.5 μg of plasmid DNA and Lipofectamine 2000 (ThermoFisher) in a 1:3 reagent:DNA ratio per well, following manufacturer’s protocol. The pGL3 plasmid was co-transfected with 20 ng of pRL-CMV to allow the normalization against renilla luciferase.

For immunofluorescence, we seeded 15.000 HeLa/PAECs per well in μ-Slide 8-well chambers (IBIDI, Germany), 24 hours later we proceeded with the transfection.

We performed the Knock-Down (KD) of KLF4, PPARγ, and VDR using a commercial pool of small interfering RNAs (siRNAs) (Dharmacon) at a concentration of 100 nM in both PAEC and HeLa cells, transfection was carried out with lipofectamine RNAiMax (ThermoFisher) following manufacturer’s protocol, 24 hours after the transfection we changed the media, and 24 hours later cells were harvested to assess KD efficiency and *EDN1* mRNA levels.

### Luciferase assay

We transfected the siRNAs in HeLa cells for 24 hours, then we changed the media, and 24 hours later we transfected the cells with the different luciferase constructs depending on the target gene (pGL3-ET1prom for *KLF4* and *PPAR*_γ_, and pmirGLO-EDN1 for *VDR*). We then proceeded with a conventional luciferase assay.

Cells were harvested 36 h post-transfection. The assay was performed using the Dual-Glo Luciferase system (Promega) following the manufacturer’s protocol, the assay was read in ½ area 96-well white plates (Corning) on an EnVision 2104 (Perkin Elmer).

Data were normalized using the firefly/renilla ratio and then relativized to the most common genotype (for the triple genotype comparison) or the empty vector (for the TF binding site test).

### qPCR

RNA extraction was carried out using NZY Total RNA Isolation Kit (NZYtech) following the manufacturer’s protocol. We used 100 ng of RNA for retrotranscription using NZY M-MuLV First-Strand cDNA synthesis kit (NZYtech). Real-time quantitative PCR (qPCR) was carried out using PowerUp SYBR Green Master Mix (ThermoFisher), 1 μL of 1:10 cDNA dilution, and the primers shown in Supp. Table 3. The reaction was performed using a total volume of 15 μL in a Step-One Plus Real-Time PCR system (ThermoFisher), cycling conditions were as following: 50 °C for 2 minutes, 95 °C for 2 minutes, 40 cycles of 95 °C for 15 seconds and 30 seconds at 60 °C; followed by a melting curve. To normalize the expression of the *KLF4, VDR, PPAR*_γ_, and *EDN1*, we followed the -ΔCT method using *YWHAZ* and *ALAS1* as reference genes.

### Chromatin Immunoprecipitation – qPCR

We used approximately 1 million HeLa cells per reaction. Chromatin shearing was performed using a sonicator (Branson). We carried out the Chromatin Immunoprecipitation (ChIP) using the ChIP Kit (abcam #ab500) following manufacturer’s protocol. For the pull down step, the following antibodies and quantities were used: anti-Histone H3 antibody as positive control (abcam, #ab1791; 2.5 μg), a Rabbit-anti-Mouse-AlexaFluor488 (ThermoFisher, #A-11059; 5 μg) and an anti-PPARγ (abcam, #ab59256; 5 μg). After DNA purification, we used 2 μL of DNA for qPCR, we carried out the reaction with the EDN1-prom primers from Supp. Table 3 using Power-Up SYBR Green Master Mix (ThermoFisher). For data analysis, we used the input percent method [100*2^(Input CT-CT(IP)].

### Immunofluorescence

We cultured PAEC cells in μ-Slide 8-well chambers (IBIDI, Germany) and performed the KD experiments by adjusting the volumes. After KD, cells were washed 3 times in PBS before being fixed with 4 % formalin for 10 minutes at 37 ºC. After washing the cells six times in PBS, we proceeded to permeabilize them in PBS+BSA 1 % (w/v) containing 0.1 % Triton X (v/v). Then, we blocked them in PBS + BSA 2 % (blocking buffer) for 1 hour at room temperature. We incubated the cells with the primary antibodies overnight in blocking buffer, washed three times with blocking buffer for 5 minutes, and incubated with the secondary antibodies and DAPI (1 μg/mL) in blocking buffer for 1 hour in the dark. Finally, we washed the chambers three times for 5 minutes in PBS and mounted them in ProLong Diamond Antifade Mountant (ThermoFisher). Images were acquired using a Leica DMI6000 inverted microscope with an integrated confocal module SP5 (Leica Microsystems, Germany). The settings used for confocal imaging were maintained in the samples

The following antibodies and dilutions were used: anti-ET1 (abcam, #ab2786, 1:500), phalloidin-Alexa488 (abcam, #ab22744, 1:1000), and Alexa Fluor 594-conjugated goat anti-mouse (ThermoFisher, #A-11005, 1:1000).

### Image analysis

All the images were processed with ImageJ (v.1.8.0). We did two batches for HeLa and two for PAECs, in each we had duplicates for each treatment and we imaged two different places per well. We then selected between 15-30 whole cells from each image and quantified the mean fluorescence for each of them on the ET-1 channel. We subtracted the background for each photo. Settings were maintained in the same IBIDI chambers and all treatments were relativized to the Mocks within their slide.

### Statistical analyses

For every experiment, we first performed a Shapiro test to assess normality. Then, we proceeded to analyze the data using a Kruskal-Wallis and a Wilcoxon test with a Holm-Sidak correction. Data were analyzed using R and plotting was performed with the ggplot2^15^ and the ggpubr packages. Comparisons were considered statistically significant when p > 0.05, for multiple comparisons we used the adjusted p-value.

### ET-1 quantification cell media

We wanted to evaluate if silencing KLF4, PPARg and VDR increased the levels of ET-1 secreted to the media. We plated PAECs in passages 2-5 in 12-well plates, at 80-90 % confluence we transfected them with the previously described conditions. We also treated another 12-well plate with DMSO (1 %), the agonist of PPARg Rosiglitazone (RGZ, 10 μM) and a PPARg antagonist GW6992 (10 μM). We had a total of 4 biological replicates. After extracting the media, we centrifuged it at 14000 g for 5 minutes to get rid of cellular debris. Also, we extracted protein from each well to correct ELISA results with the total protein present in the cells.

We quantified ET-1 levels using the Endothelin-1 Quantikine ELISA kit (R&D systems). First we diluted the media 1/125 in the appropriate buffer and used 75 μL of the dilution for the assay. We incubated the media with 200 μL of assay buffer for 1 hour at RT in a shaker. Then we washed the plate and incubated with the anti-ET-1 conjugated antibody for 3 hours at RT in a shaker. Finally, we washed the plate and added the substrate. We incubated it for 30 minutes before adding the stop solution and reading the results in a plate iMark microplate absorbance reader (BioRad).

## Results

### Common variation can be found in EDN1 regulatory regions

Sanger sequencing of *EDN1* was carried out in patients with IPAH. Variants were only found in the regulatory regions of the gene, not in the coding region, which is highly conserved. In the 5’UTR we found c.-131delA (rs397751713; Figure 1A) and in the 3’ regulatory region g.12298751G>A (rs2859338; Figure 1A). Both of them are common SNPs in the European population and are classified as Benign in Varsome. Although we found slight differences in the frequencies (Table 1), they did not meet statistical significance. After a closer *in silico* analysis we decided to carry out functional assays.

**Figure 1.**
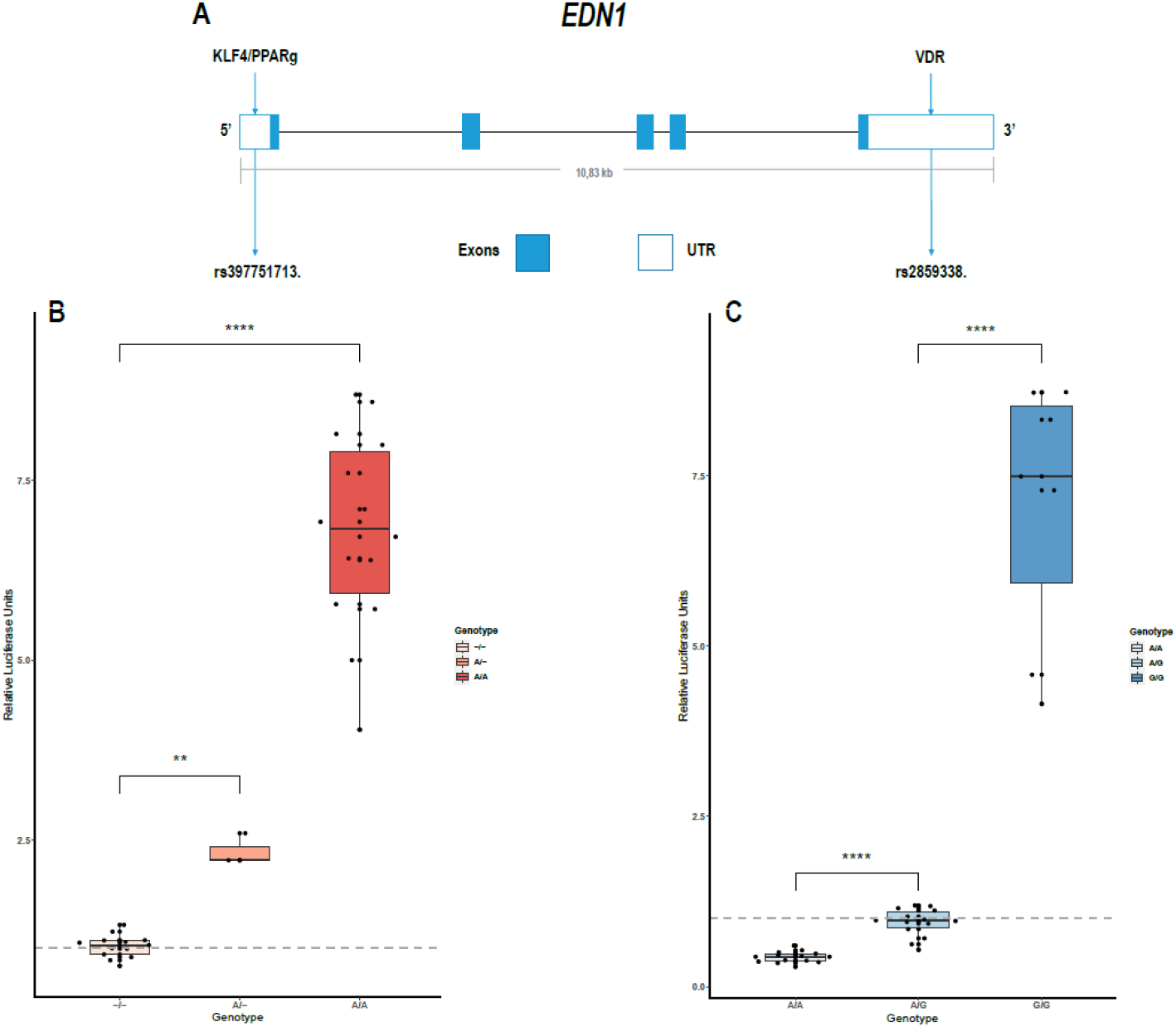
Common variants in *EDN1* UTRs show different activity depending on the genotype. A) Schematic view of the *EDN1* gene with the location of the SNPs and the pre transcription factors to interact with them. B) Comparison of the different genotypes of rs397751713 show increased activity in the A/A genotype and A/- when compared to the comm genotype -/- (n = 2 for A/-; n = 6 the rest). C) Comparison of the different genotypes of rs2859338 show increased G/G genotype activity and reduced activity in A/A compared wi1 commonest genotype A/G (n = 4). Data are represented as box plots depicting the quartiles (** p > 0.01, **** p > 0.0001).

**Table 1.** Frequencies of the SNPs detected in *EDN1* UTRs in our cohort and ensemble.

| **rs397751713** |  |  |  |
| --- | --- | --- | --- |
| **Genotype** | **Patients** | **Frequency** | **Ensembl** |
| A/A | 3 | 0.08 | 0.112 |
| A/- | 10 | 0.28 | 0.364 |
| -/- | 23 | 0.64 | 0.523 |
| **rs2859338** |  |  |  |
| G/G | 10 | 0.30 | 0.355 |
| G/A | 20 | 0.61 | 0.514 |
| A/A | 3 | 0.09 | 0.131 |

### The common SNPs are predicted to lower the affinity of some transcription factors

We used the bioinformatics software Genomatix to predict the transcription factors binding in the whole 5’UTR and 3’ regulatory region. We did this step twice changing only the SNPs. Among the 65 possible candidates provided, those with binding sites that included the SNPs detected were selected. This way, the candidates were reduced to KLF4 (Krüppel-Like Factor 4), PPARγ (Peroxisome proliferator-activated receptor _γ_) and HNF4A (nuclear hepatocyte factor 4_α_) for 5’UTR and VDR (Vitamin D receptor) in the 3’ position.

Within these four candidates, we were only able to establish a nexus with PAH in three cases: KLF4, PPARγ, and VDR, with values for its similarity matrix of 0.961, 0.842, and 0.974 respectively.

### Promoters carrying the different *EDN1* c.-131del (rs397751713) genotypes show differential activity *in vitro*

We cloned into the p.GL3 *EDN1* promoters carrying the different genotypes of the common SNP rs397751713. We found that the most common form c.-131del in homozygosity shows lower expression (-/-; 1.02 ± 0.15) when compared with the less common ancestral adenine insertion, both in heterozygosity (A/-; 2.35 ± 0.21; p > 0.01) and homozygosity (A/A; 6.79 ± 1.36; p > 0.0001) (Figure 1B).

### The 3’ regulatory region variant g.12298751G>A (rs2859338) genotypes show different activity *in vitro*

In the same manner, we tested the effect of the different genotypes of rs2859338 using p.mirGLO as this SNP is located in the 3’ of *EDN1*. In this case, we used the most common heterozygous form to relativize and compare (A/G; 0.94 ± 0.2). The homozygous genotype A/A showed less expression than the most common variant (A/A; 0.43 ± 0.08; p > 0.0001), while the less common A>G substitution in homozygosity showed a high increase in luciferase expression (G/G; 7.04 ± 1.91; p > 0.0001) (Figure 1C).

### The KD of KLF4, PPARγ, and VDR increase *EDN1* mRNA levels in PAECs

We measured *EDN1* mRNA production 48 hours after the KD of the KLF4, PPARγ, and VDR. We found the overall *EDN1* levels increased in all of them when compared to the Mock siRNA (Figure 2). siPPARγ showed the greatest increase (1.31 ± 0.17, p > 0.0001), followed by siKLF4 (1.18 ± 0.08, p > 0.001) and then siVDR (1.1 ± 0.09, p > 0.02).

**Figure 2.**
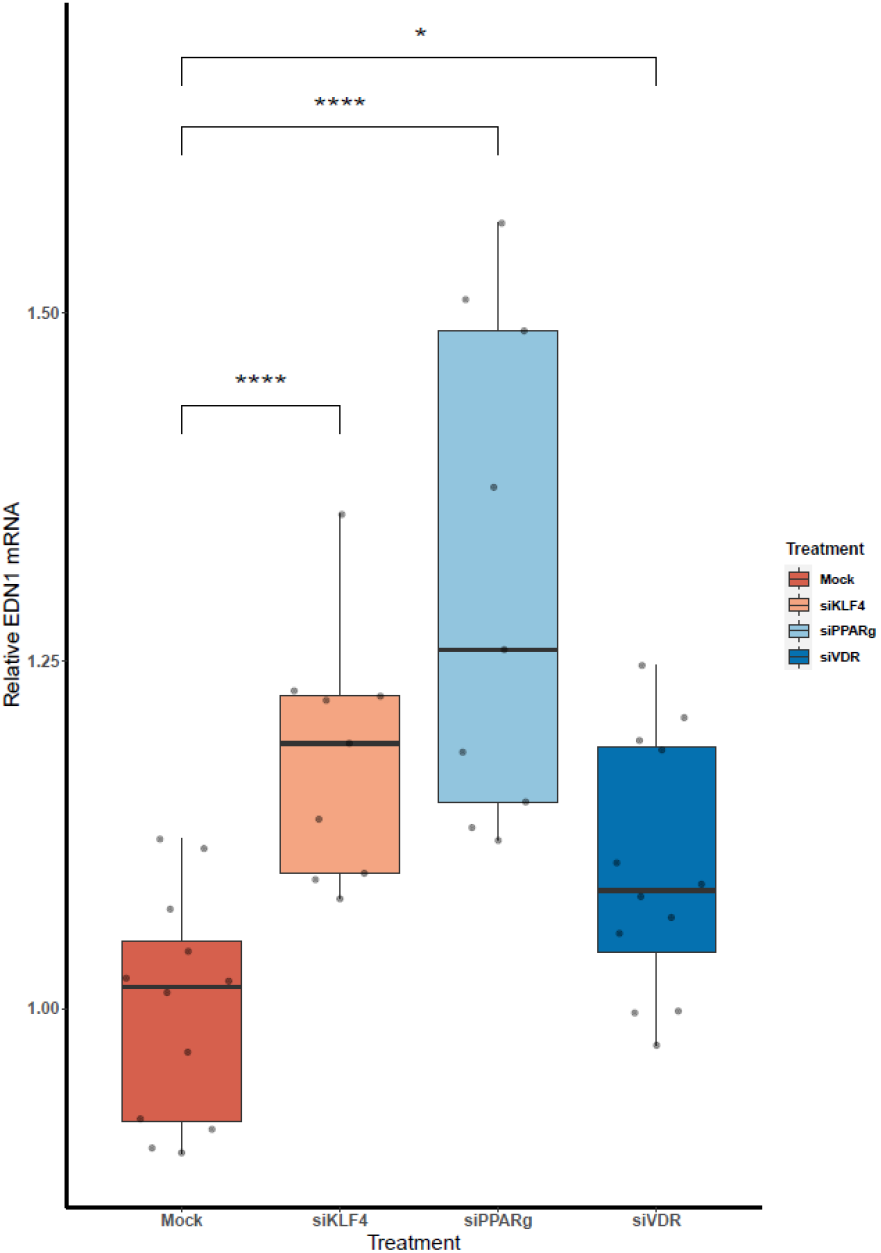
The KD of PPARγ, KLF4, and VDR increase the mRNA levels of *EDN1*. (n = 3) Data are shown as box plots depicting the quartiles (* p > 0.05, **** p > 0.0001).

### PPARγ binds to the EDN1 promoter and is influenced by the A/A genotype of rs397751713

To test if rs397751713 genotypes influenced the binding of KLF4 or PPARγ, we did a KD of KLF4 using siRNA. We first optimized the reaction in primary PAECs getting only around a 38 % inhibition for KLF4 (Figure 3A), while for PPARγ it went up to 98 % (Figure 3B), in both cases we used the maximum recommended siRNA amount.

**Figure 3.**
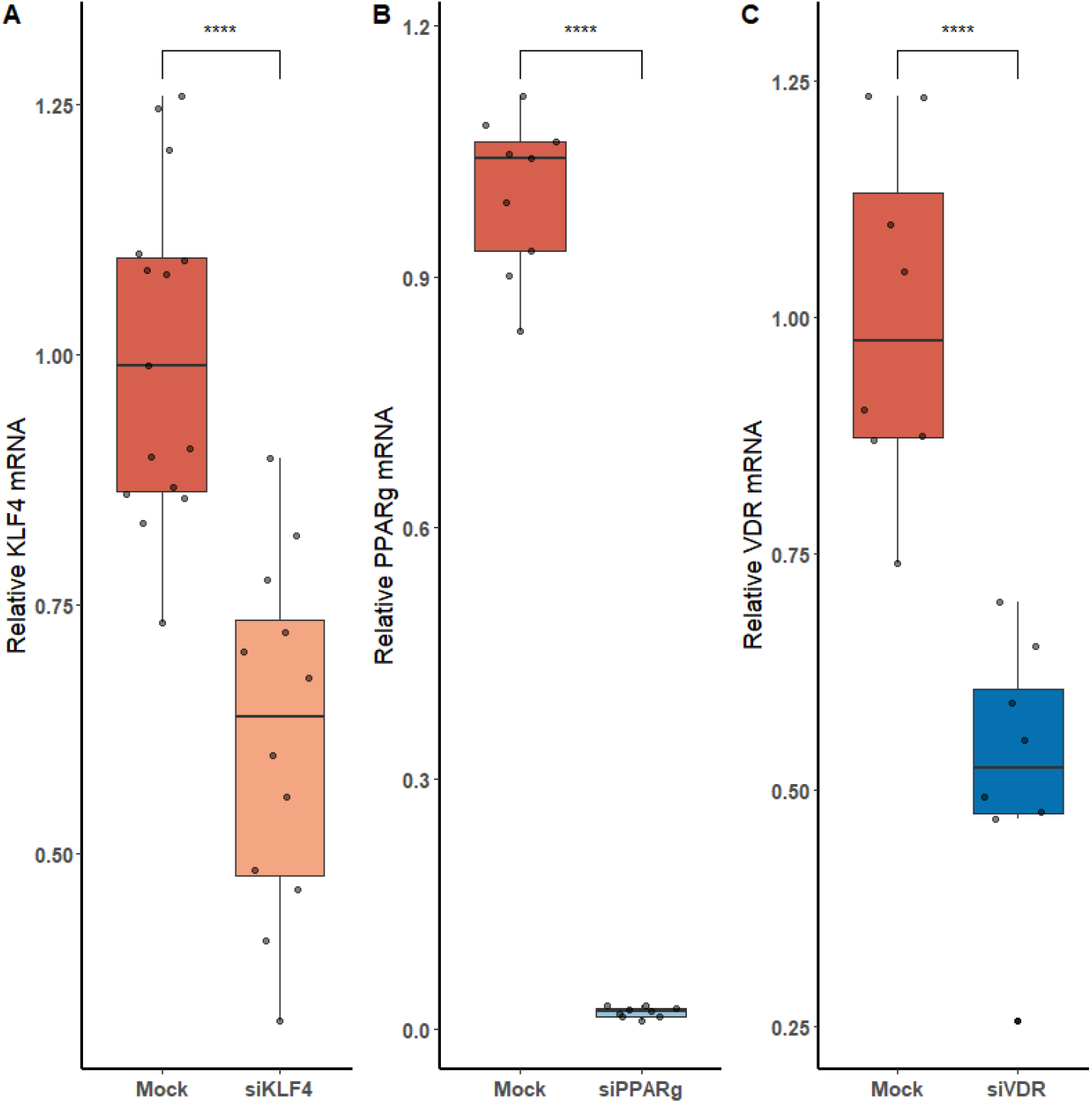
Knock-Down efficiency for KLF4, PPARγ and VDR. Data are shown as box plots representing median ± quartiles. Dots represent biological replicates.

Then, we replicated the conditions in HeLa and transfected the p.GL3-EDN1prom with the homozygous genotypes (-/- and A/A). The results showed the same pattern between mock and siKLF4 (Figure 4A). However, siPPARγ showed a completely different pattern compared to control (Fig. 3A), increasing luciferase activity to similar levels between -/- and A/A (7.96 ± 0.93 vs 7.06 ± 0.35). This indicates that PPARγ binds in this position and its affinity could be reduced by the A/A genotype.

**Figure 4.**
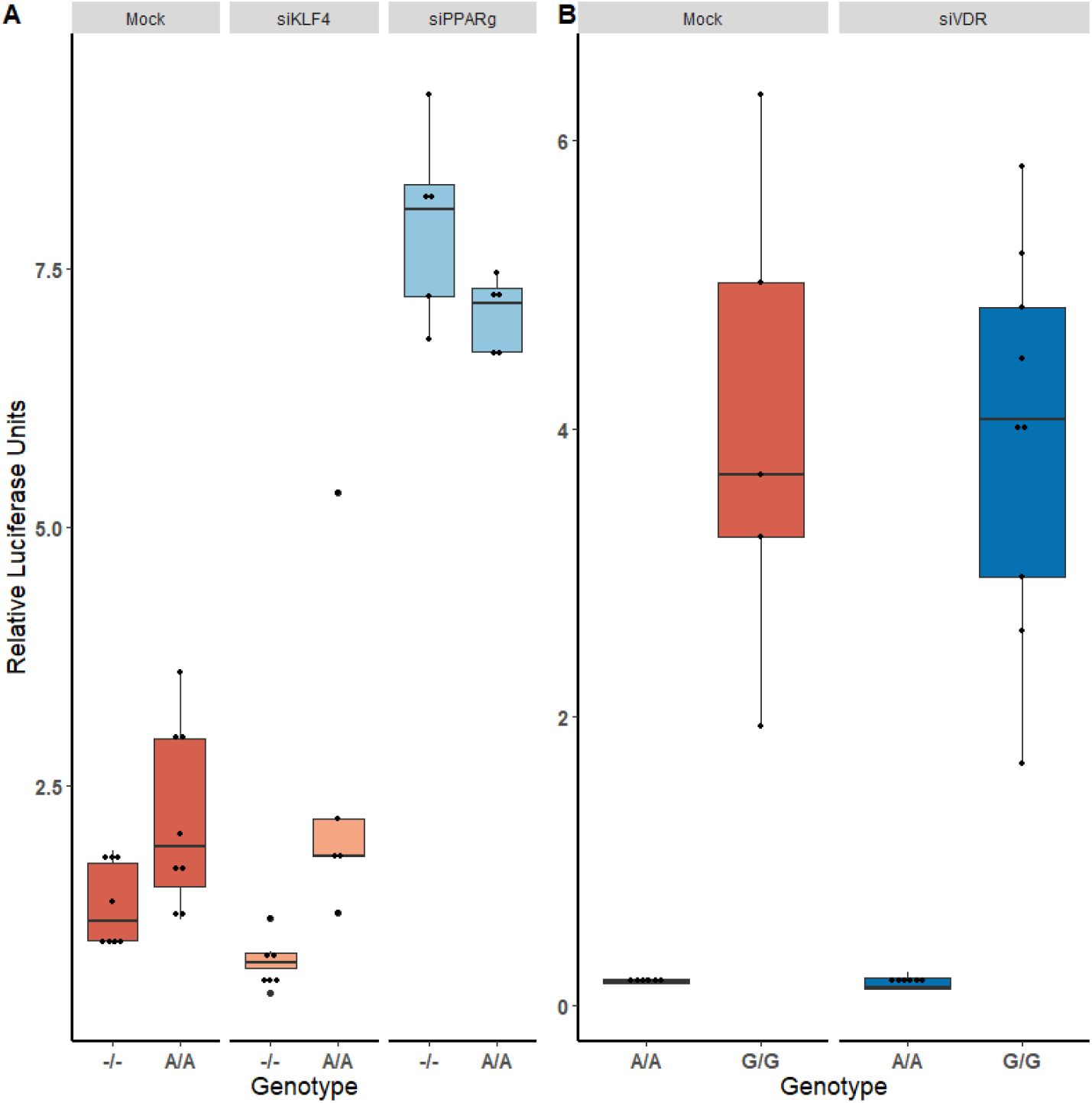
KD experiments demonstrate that PPARγ binds in rs397751713. A) Comparison of the luciferase activity in the homozygous genotypes of rs397751713 after different treatments, siKLF4 and Mock show the same pattern while siPPARγ increases the activity of both genotypes to similar levels (n =. B) Comparison of the luciferase activity in the homozygous genotypes of rs2859338 after Mock or siVDR treatment shows the same pattern. Data are shown as box plots representing median ± quartiles. Dots represent biological replicates.

In the case of rs2859338, we silenced VDR to test if it was bound in our target region. After optimization, we managed to lower VDR expression by only 48 % (Figure 3C). The luciferase results showed the same pattern for both genotypes treated with Mock (A/A 0.16 ± 0.01; G/G 4.04 ± 1.68) and siVDR (A/A 0.15 ± 0.05; G/G 3.96 ± 1.33) (Figure 4B). Also, we were able to amplify this region after carrying out a ChIP-qPCR assay, (Figure 5).

**Figure 5.**
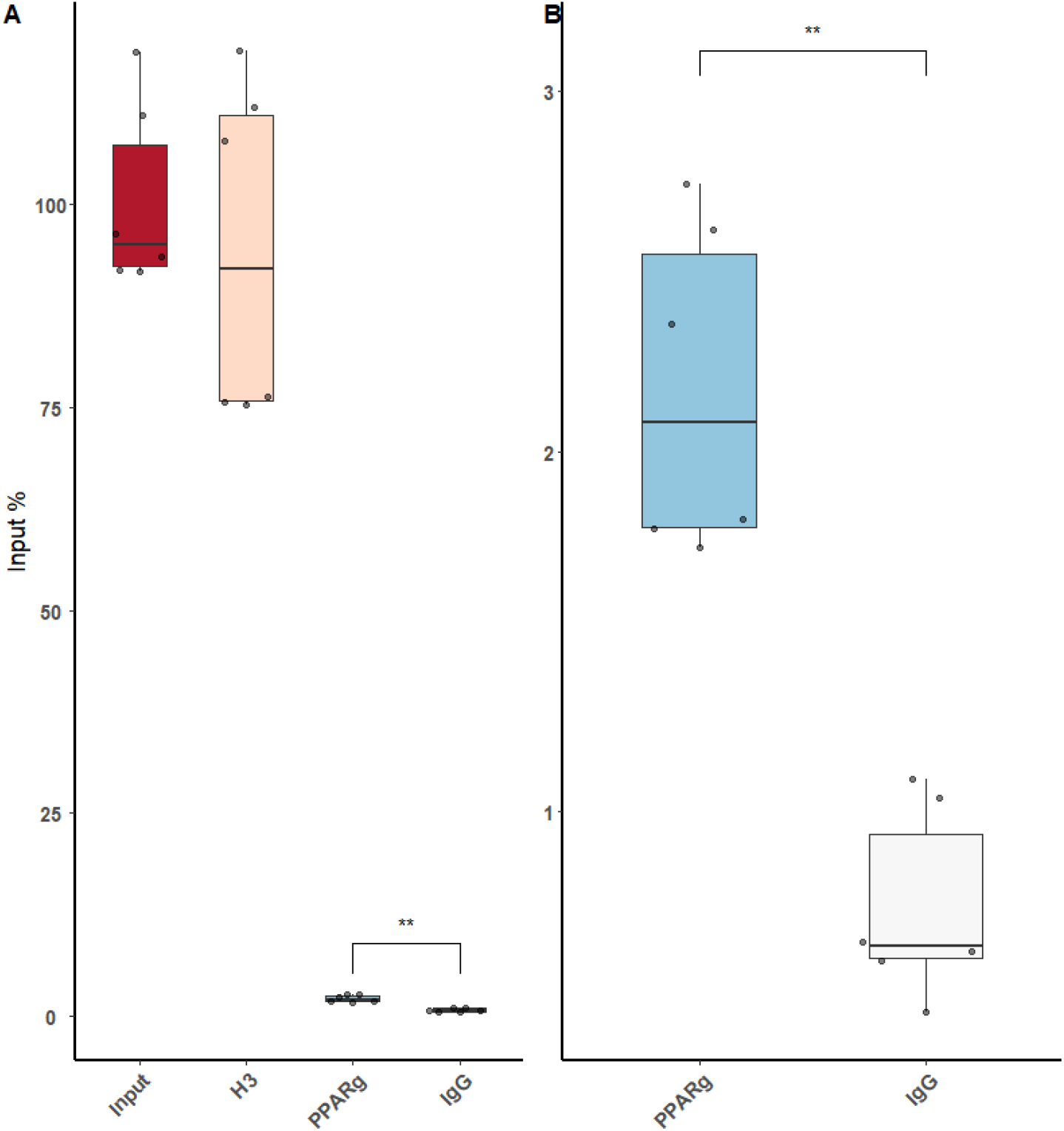
ChIP-qPCR using an anti-PPARγ antibody demonstrates that low amounts of PPARγ binds around rs397751713. A) Full plot with the input chromatin used in the assay, a positive control (H3) and a Rabbit IgG. B) Comparison between the control IgG and PPARγ amplification.

### PPARγ KD increases ET-1 production in HeLa and PAECs, while KLF4 does it only in HeLa and VDR in PAECs

We used immunofluorescence to confirm that the increase at mRNA levels led to higher amounts of ET-1 production in PAECs. We first experimented in HeLa cells to see if a greater KD efficiency for VDR and KLF4 changed the results. HeLa cells showed increase ET-1 levels after the treatment with siKLF4 (1.31 ± 0.32, p > 0.0001) and siPPARγ (1.68 ± 0.43, p > 0.0001) while siVDR had more or less the same as WT (0.94 ± 0.23, n.s.) (Figure 6A and 6B). In PAECs, siPPARγ showed the highest increase in ET-1 (1.57 ± 0.66, p > 0.0001) followed by siVDR (1.55 ± 0.53, p > 0.0001), siKLF4 had a slight increase barely significant (1.17 ± 0.45, p > 0.031) (Figure 6C and 6D).

**Figure 6.**
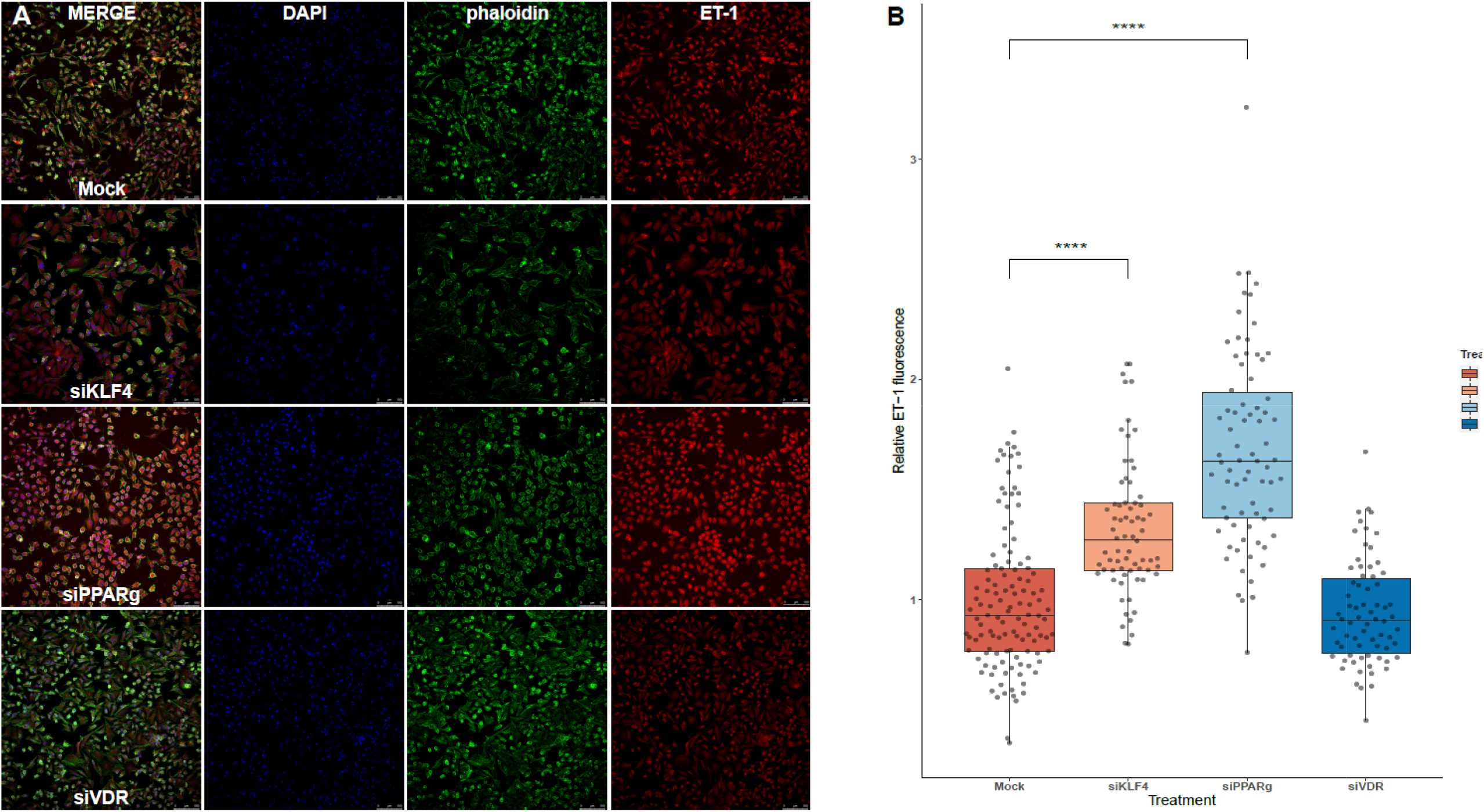
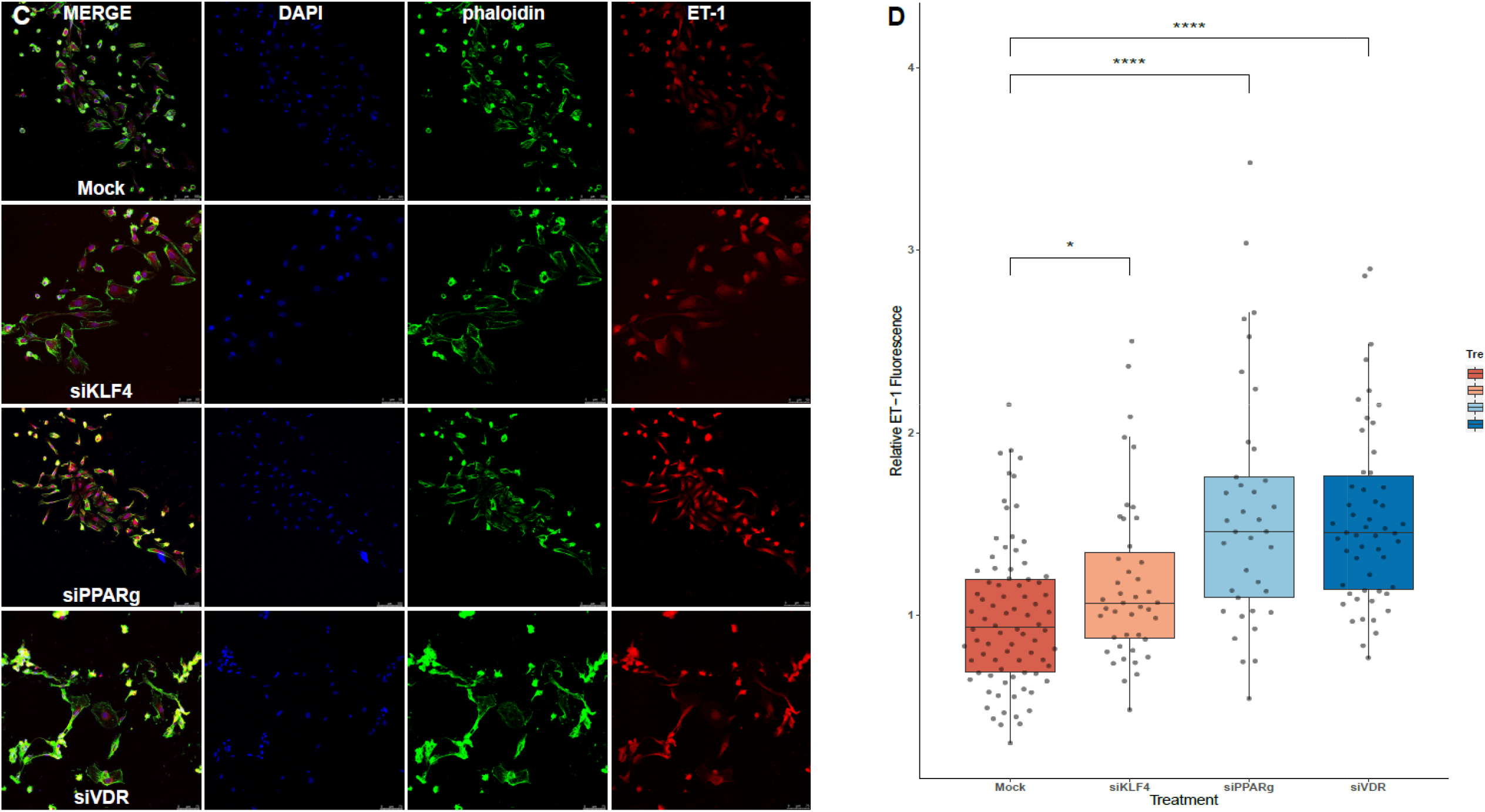
Silencing PPARγ increases ET-1 production in HeLa and PAECs while VDR does only in PAECs and KLF4 in HeLa. A) Representative immunofluorescences for HeLa ce Quantification of ET-1 production using immunofluorescence in HeLa, treatment with siKLF4 and siPPARγ increases ET-1 fluorescence (n= 4). C) Representative immunofluorescenc PAECs. D) Quantification of ET-1 production using immunofluorescence in PAECs, treatment with siPPARγ and siVDR show the highest increase in ET-1 fluorescence, while siKLF4 has a increase (n = 6). Data are represented as box plots depicting the quartiles (* p > 0.05, **** p > 0.0001).

### ET-1 levels in media are not increased after silencing

After running an ELISA against ET-1 using cell-culture media, we did not find any statistically significant difference between the treatments. Also, attenuating or activating PPARγ did not change the overall levels of ET-1 in the media (Figure 7).

**Figure 7.**
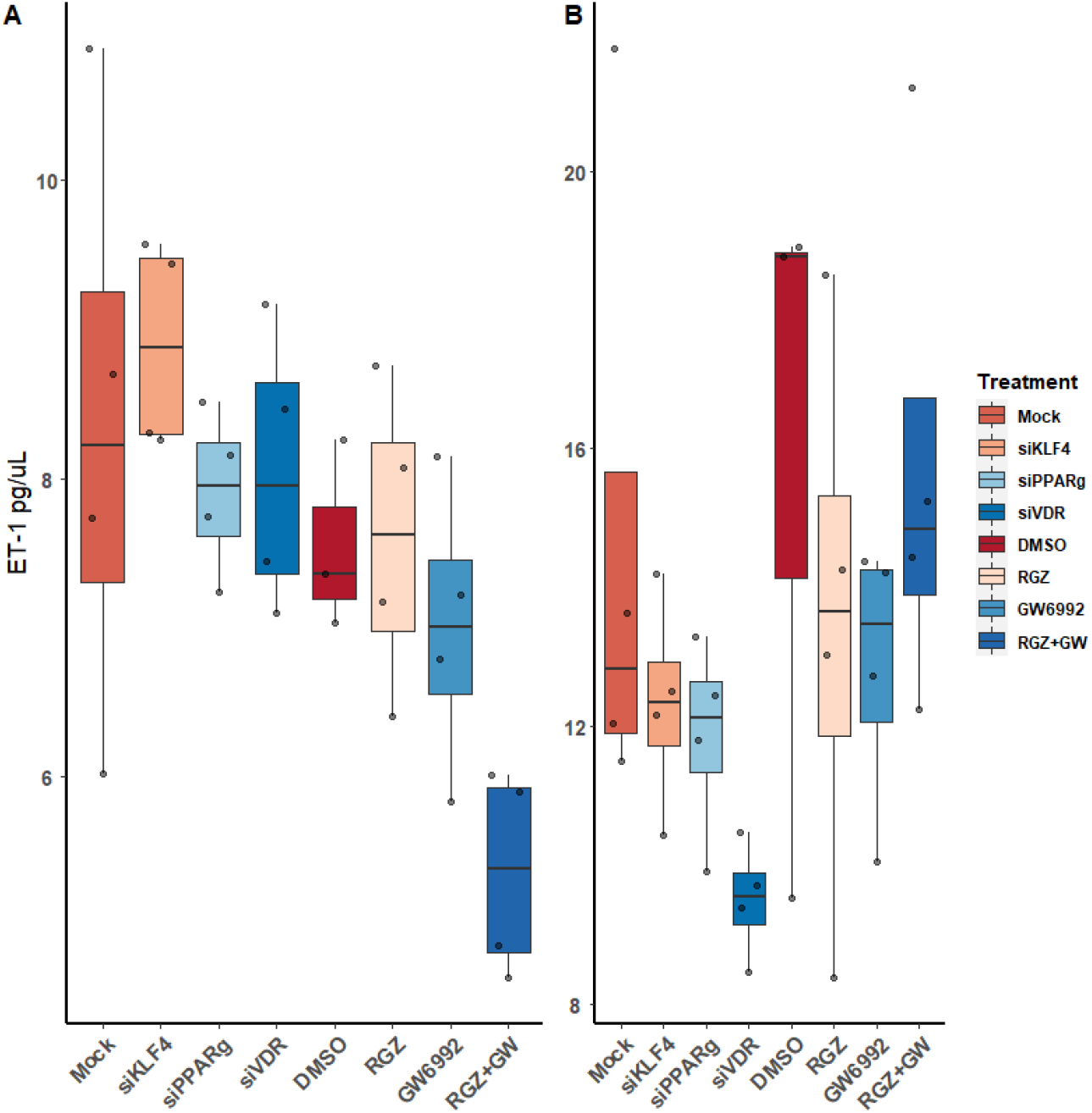
Quantification of ET-1 by sandwich ELISA. A) Raw values. B) After normalizing against the total protein from the well. Data are shown as box plots representing median ± quartiles. Dots represent biological replicates.

## Discussion

Genetic screening in PAH is usually performed by limited gene panels or whole-exome sequencing that neglect non-coding regions. Regulatory regions as promoters, UTRs and other regulatory regions have been historically understudied.

In this study, we analyzed *EDN1* regulatory regions in the search for variants that could modulate *EDN1* expression, and then we coupled functional studies with patients’ data to dissect the effect of these changes in a translational way.

In our screening, we only found three variants, the two common SNPs rs397751713 and rs2859338 in regulatory regions, and the already reported p.Lys198Asn within the coding sequence^12^. The luciferase assays we performed for these non-coding variants showed a marked increase in the luciferase activity for the less frequent homozygous genotype of both SNPs. Looking at this data from an evolutionary perspective, we realized that the less frequent genotypes of both these SNPs were the ancestral forms, still present in the oldest clades. Besides, ET-1 is a very well conserved and potent vasoconstrictor^15^. It would make sense that evolution had fixed the variations that helped regulate its expression.

Using *in silico* tools we predicted that KLF4/PPARγ could bind right where rs397751713 is located, while VDR did it right in rs2859338. We were able to determine that silencing PPARγ increased the luciferase levels in all the tested genotypes, while the silencing of KLF4 showed the same pattern as the Mock. This suggests that PPARγ bind at this position, and at least in our static conditions KLF4 is not. PPARγ seems to interact with both genotypes, so the expression differences between the genotypes could be a matter of affinity, as PPARγ may be able to bind more efficiently to -/- genotype leading to an inhibition of the *EDN1* expression, while the Adenine insertion would have less PPARγ binding affinity and thus, a higher expression of *EDN1*. However, silencing VDR showed the same results as in the basal luciferase assay, meaning that VDR cannot bind at rs2859338.

To evaluate the influence that these TFs play on ET-1 levels, we performed KD experiments in PAECs. We first measured *EDN1* mRNA and found increased levels in all siRNA treatments. The result was almost non-significant in the case of siKLF4 with barely a 10 % increase, while siVDR showed an 18 % increase, and siPPARγ a 31 %. Using the same conditions, we used immunofluorescence to measure ET-1 protein production at a cellular level, first in HeLa, and then in PAECs. Both of these cell lines showed increased ET-1 levels of more than 50 % after silencing PPARγ. While silencing KLF4 was more effective in HeLa (31 %) than it did in PAECs (17 %), however, VDR only resulted in increased ET-1 production in PAECs (55 %). Altogether, in a static culture, PPARγ probably plays a bigger role in regulating ET-1 than KLF4, as its expression is triggered by shear stress^16^, or VDR do, as maybe their influence could be played indirectly.

KLF4 is a well-known TF in the PAH context. It is expressed in the vascular endothelium, promoting anti-inflammatory and anticoagulant states. The lack of this transcription factor in the vascular endothelium was shown to exacerbate hypoxia-induced PAH and increase the expression of ET-1^17^. Our results support this, and recent ChIP-seq data show that it binds to the *EDN1* promoter in the position we studied^16^.

Besides, PPARγ influence in ET-1 mediated vascular damage is well-known^18^, but how this interaction happened has not been shown until now. PPARγ is a ligand-dependent TF, which binds to hormonal response elements in promoters of target genes, mainly related to adipogenesis and secondarily to glucose metabolism^19^. In PAH, remodeled and muscularized precapillary arterioles show frequently reduced PPARγ expression in endothelial cells. PPARγ not only regulates hypoxia-induced ET-1 levels but also other components of the ET-1 signaling pathway, such as ECE-1 mRNA levels, ETA, and ETB^20^. Due to the above, it is believed that PPARγ agonists could reverse pulmonary vascular remodeling^21,22^. Furthermore, a malfunction of BMPR2 has been shown to decrease endogenous PPARγ activity and promote metabolic pathways associated with vascular remodeling^23^. Experiments in animal models support the results obtained in this work^18^. Furthermore, it appears that there is a reciprocal regulation between Endothelin and PPARγ ^24^.

The notion that several extrapulmonary organs (heart, skeletal muscle, and adipose tissue) show vascular and metabolic abnormalities suggests that PAH is a systemic rather than exclusively pulmonary hypertensive disease. Dyslipidemia and insulin resistance are evident in animal models of PAH and human disease^25^. In fact, many drugs that act as ligands for the PPARγ receptor are currently used as a treatment for type 2 diabetes^20^.

The vitamin D receptor (VDR) is a transcription factor that is activated in the presence of calcitriol, the active form of vitamin D. The relationship of VDR with genes involved in the regulation of the vascular tone, such as ET-1 (vasoconstrictor) and nitric oxide (NO, vasodilator), has been demonstrated. The regulation of NO by the VDR is direct, but it does not appear to be so in the case of ET-1. Previous reports showed that the regulation of ET-1 by VDR would not alter the expression of preproendothelin, but would act on the Endothelin-Converting Enzyme 1 (ECE1), generating more active endothelin^27^. However, VDR increases the amount of NO by directly activating eNOS, generating a vasodilator effect^26^. So it would be expected that the system would saturate rapidly in the case of ET-1, as the amount of preproendothelin mRNA did not increase^26^, and the overall result would be vasodilation. This hypothesis is supported by a recent report by Callejo *et al*.^27^, and our results. As after silencing *VDR*, ET-1 expression increased *in vitro*.

ELISA technique demonstrated that even though ET-1 production is increased at cell level, it does not increase the levels of ET-1 secreted to cell media. None of the treatments tested showed any relevant difference when compared with the controls, and even pharmacological inhibition or activation of PPARγ had no effect in the secretion of ET-1. This lead us to think that ET-1 secretion may be tightly regulated and could be stopped after reaching a certain level of ET-1 in the media, it could also be recaptured by ETBR in the ECs, or it could be affected by the absence of the SMCs that would usually be the target of this peptide. Our fluorescence data show the accumulation of vesicles around the nuclei, but not near the membrane to be secreted^28^, which make sense with what we detected in the ELISA.

PAH is a complex disease where genetic information is limited to the main causal genes and little is known about the role of common variation^29,30^. Over the last decades, the capability of detecting genetic variation has increased; we can now try to use common variation to explain phenotypic expressivity. Common variants influencing the regulation of the ET-1 pathway had been proposed previously when it was shown that the response of PAH patients to ERAs could be modified by a common intronic SNP in the GNG2 gene^31^. But in that study, no SNP in EDN1 was statistically significant or studied in-depth, and the molecular effect of the GNG2 SNP was not analyzed *in vitro*. Another example in a closely related pathway is the Angiotensin II type 1 receptor (AGTR1), where patients harboring a homozygous C/C allele for rs5186 showed a later age of diagnosis^32,33^. Also, a SNP (rs12483377) in the Endostatin gene (Col18a1) showed different serum levels in patients that were homozygous for p.Asp1675Asn instead of the ancestral Asparagine, while the levels of Endostatin impacted survival^34^. But common variants influencing PAH not only appear in cardiovascular-related pathways, a SNP in *Sirtuin 3* (*SIRT3*; rs11246020) was associated with IPAH, as it lowered SIRT3 activity a 30 % favoring glycolysis in mitochondria^35^. The latest additions to this list are a SNP in an enhancer locus near SOX17 (rs10103692) that was associated with PAH, and a variant in *HLA-DPA1/DPB1* (rs2856830) that was associated with I/HPAH and showed different survival depending on the genotype (CC vs TT)^36^.

The mutational load could modulate gene expression and alter patients’ phenotypes. PAH variability could be explained by how common variation influences the different pathways involved in the pathogenesis, it could be a way to explain how mutations with the same effect can have very different phenotypes. The next step should be screening the SNPs we identified in patients undertaking ERAs, as they could explain the very different response to this treatment. We would like to encourage the biggest cohorts to go beyond rare variation and test in detail possible genetic modulators.

The main limitations of this study are that due to the small size of our cohort we cannot draw conclusions at the different outcome levels. Moreover, we have been working with plasmids constructed with limited fragments of the regulatory regions of *EDN1* in cell lines. The results on KLF4 and VDR could need a better silencing efficiency to show in the luciferase assay, our confidence in them is based on repetition and that we can see the same patterns at mRNA and protein level. Besides, culturing PAECs in physiological conditions could have changed some of our results.

In conclusion, we show how common variants in *EDN1* regulatory regions could alter ET-1 levels. We validated that PPARγ binds in rs397751713 and heavily influences ET-1 regulation *in vitro*. Furthermore, KLF4 and VDR influence ET-1 production in a cell-dependent manner.

## Acknowledgements

We thank Sebastián Comesaña, Verónica Outeiriño and Inés Pazos of the *Centro de Apoio Científico-Técnico á Investigación* (CACTI) for their help in sequencing and imaging. We thank Ana Paula Borges Diez for her help with Illustrator.

## References

1. Galiè N, Humbert M, Vachiery J-L, Gibbs S, Lang I, Torbicki A, Simonneau G, Peacock A, Vonk Noordegraaf A, Beghetti M, Ghofrani A, Gomez Sanchez MA, Hansmann G, Klepetko W, Lancellotti P, Matucci M, McDonagh T, Pierard LA, Trindade PT, Zompatori M, Hoeper M. 2015 ESC/ERS Guidelines for the diagnosis and treatment of pulmonary hypertension. European Respiratory Journal 2015;46:903–975.

2. Morrell NW, Aldred MA, Chung WK, Elliott CG, Nichols WC, Soubrier F, Trembath RC, Loyd JE. Genetics and genomics of pulmonary arterial hypertension. European Respiratory Journal 2018;1801899.

3. Galiè N, Channick RN, Frantz RP, Grünig E, Jing ZC, Moiseeva O, Preston IR, Pulido T, Safdar Z, Tamura Y, McLaughlin VV. Risk stratification and medical therapy of pulmonary arterial hypertension. Eur Respir J 2019;53.

4. Inoue A, Yanagisawa M, Kimura S, Kasuya Y, Miyauchi T, Goto K, Masaki T. The human endothelin family: three structurally and pharmacologically distinct isopeptides predicted by three separate genes. Proc Natl Acad Sci U S A 1989;86:2863–2867.

5. Dupuis J, Stewart DJ, Cernacek P, Gosselin G. Human pulmonary circulation is an important site for both clearance and production of endothelin-1. Circulation 1996;94:1578–1584.

6. Davenport AP, Hyndman KA, Dhaun N, Southan C, Kohan DE, Pollock JS, Pollock DM, Webb DJ, Maguire JJ. Endothelin. Pharmacol Rev 2016;68:357–418.

7. Rubens C, Ewert R, Halank M, Wensel R, Orzechowski HD, Schultheiss HP, Hoeffken G. Big endothelin-1 and endothelin-1 plasma levels are correlated with the severity of primary pulmonary hypertension. Chest 2001;120:1562–1569.

8. Montani D, Souza R, Binkert C, Fischli W, Simonneau G, Clozel M, Humbert M. Endothelin-1/endothelin-3 ratio: a potential prognostic factor of pulmonary arterial hypertension. Chest 2007;131:101–108.

9. Latus H, Karanatsios G, Basan U, Salser K, Müller S, Khalil M, Kreuder J, Schranz D, Apitz C. Clinical and prognostic value of endothelin-1 and big endothelin-1 expression in children with pulmonary hypertension. Heart 2016;102:1052–1058.

10. Giaid A, Yanagisawa M, Langleben D, Michel RP, Levy R, Shennib H, Kimura S, Masaki T, Duguid WP, Stewart DJ. Expression of endothelin-1 in the lungs of patients with pulmonary hypertension. N Engl J Med 1993;328:1732–1739.

11. Satwiko MG, Ikeda K, Nakayama K, Yagi K, Hocher B, Hirata K, Emoto N. Targeted activation of endothelin-1 exacerbates hypoxia-induced pulmonary hypertension. Biochem Biophys Res Commun 2015;465:356–362.

12. Pousada G, Baloira A, Vilariño C, Valverde D. [K198N polymorphism in the EDN1 gene in patients with pulmonary arterial hypertension]. Med Clin (Barc) 2015;144:348–352.

13. Cartharius K, Frech K, Grote K, Klocke B, Haltmeier M, Klingenhoff A, Frisch M, Bayerlein M, Werner T. MatInspector and beyond: promoter analysis based on transcription factor binding sites. Bioinformatics 2005;21:2933–2942.

14. Kent WJ, Sugnet CW, Furey TS, Roskin KM, Pringle TH, Zahler AM, Haussler D. The human genome browser at UCSC. Genome research Cold Spring Harbor Laboratory Press; 2002;12:996–1006.

15. Square TA, Jandzik D, Massey JL, Romášek M, Stein HP, Hansen AW, Purkayastha A, Cattell MV, Medeiros DM. Evolution of the endothelin pathway drove neural crest cell diversification. Nature 2020;585:563–568.

16. Moonen J-RAJ, Chappell J, Shi M, Shinohara T, Li D, Mumbach MR, Zhang F, Nasser J, Mai DH, Taylor S, Wang L, Metzger RJ, Chang HY, Engreitz JM, Snyder MP, Rabinovitch M. KLF4 Recruits SWI/SNF to Increase Chromatin Accessibility and Reprogram the Endothelial Enhancer Landscape under Laminar Shear Stress | bioRxiv. bioRxiv.

17. Shatat MA, Tian H, Zhang R, Tandon G, Hale A, Fritz JS, Zhou G, Martínez-González J, Rodríguez C, Champion HC, Jain MK, Hamik A. Endothelial Krüppel-Like Factor 4 Modulates Pulmonary Arterial Hypertension. Am J Respir Cell Mol Biol 2014;50:647–653.

18. Idris-Khodja N, Ouerd S, Trindade M, Gornitsky J, Rehman A, Barhoumi T, Offermanns S, Gonzalez FJ, Neves MF, Paradis P, Schiffrin EL. Vascular smooth muscle cell peroxisome proliferator-activated receptor _γ_ protects against endothelin-1-induced oxidative stress and inflammation. J Hypertens 2017;35:1390–1401.

19. Floyd ZE, Stephens JM. Controlling a master switch of adipocyte development and insulin sensitivity: covalent modifications of PPARγ. Biochim Biophys Acta 2012;1822:1090–1095.

20. Kang B-Y, Kleinhenz JM, Murphy TC, Hart CM. The PPARγ ligand rosiglitazone attenuates hypoxia-induced endothelin signaling in vitro and in vivo. Am J Physiol Lung Cell Mol Physiol 2011;301:L881–891.

21. Liu Y, Tian XY, Huang Y, Wang N. Rosiglitazone Attenuated Endothelin-1-Induced Vasoconstriction of Pulmonary Arteries in the Rat Model of Pulmonary Arterial Hypertension via Differential Regulation of ET-1 Receptors. PPAR Res 2014;2014:374075.

22. Palacios-Ramírez R, Hernanz R, Martín A, Pérez-Girón JV, Barrús MT, González-Carnicero Z, Aguado A, Jaisser F, Briones AM, Salaices M, Alonso MJ. Pioglitazone Modulates the Vascular Contractility in Hypertension by Interference with ET-1 Pathway. Sci Rep 2019;9:16461.

23. Hansmann G, Wagner RA, Schellong S, Perez VA de J, Urashima T, Wang L, Sheikh AY, Suen RS, Stewart DJ, Rabinovitch M. Pulmonary arterial hypertension is linked to insulin resistance and reversed by peroxisome proliferator-activated receptor-gamma activation. Circulation 2007;115:1275–1284.

24. Wolf D, Tseng N, Seedorf G, Roe G, Abman SH, Gien J. Endothelin-1 decreases endothelial PPARγ signaling and impairs angiogenesis after chronic intrauterine pulmonary hypertension. Am J Physiol Lung Cell Mol Physiol 2014;306:L361–371.

25. Legchenko E, Chouvarine P, Borchert P, Fernandez-Gonzalez A, Snay E, Meier M, Maegel L, Mitsialis SA, Rog-Zielinska EA, Kourembanas S, Jonigk D, Hansmann G. PPARγ agonist pioglitazone reverses pulmonary hypertension and prevents right heart failure via fatty acid oxidation. Sci Transl Med 2018;10.

26. Martínez-Miguel P, Valdivielso JM, Medrano-Andrés D, Román-García P, Cano-Peñalver JL, Rodríguez-Puyol M, Rodríguez-Puyol D, López-Ongil S. The active form of vitamin D, calcitriol, induces a complex dual upregulation of endothelin and nitric oxide in cultured endothelial cells. Am J Physiol Endocrinol Metab 2014;307:E1085–1096.

27. Callejo M, Mondejar-Parreño G, Morales-Cano D, Barreira B, Esquivel-Ruiz S, Olivencia MA, Manaud G, Perros F, Duarte J, Moreno L, Cogolludo A, Perez-Vizcaíno F. Vitamin D deficiency downregulates TASK-1 channels and induces pulmonary vascular dysfunction. Am J Physiol Lung Cell Mol Physiol 2020;319:L627–L640.

28. Stow LR, Jacobs ME, Wingo CS, Cain BD. Endothelin-1 gene regulation. FASEB J 2011;25:16–28.

29. Southgate L, Machado RD, Gräf S, Morrell NW. Molecular genetic framework underlying pulmonary arterial hypertension. Nature Reviews Cardiology Springer Science and Business Media LLC; 2019;

30. Swietlik EM, Prapa M, Martin JM, Pandya D, Auckland K, Morrell NW, Gräf S. ‘There and Back Again’-Forward Genetics and Reverse Phenotyping in Pulmonary Arterial Hypertension. Genes (Basel) 2020;11.

31. Benza RL, Gomberg-Maitland M, Demarco T, Frost AE, Torbicki A, Langleben D, Pulido T, Correa-Jaque P, Passineau MJ, Wiener HW, Tamari M, Hirota T, Kubo M, Tiwari HK. Endothelin-1 Pathway Polymorphisms and Outcomes in Pulmonary Arterial Hypertension. Am J Respir Crit Care Med 2015;192:1345–1354.

32. Chung WK, Deng L, Carroll JS, Mallory N, Diamond B, Rosenzweig EB, Barst RJ, Morse JH. Polymorphism in the angiotensin II type 1 receptor (AGTR1) is associated with age at diagnosis in pulmonary arterial hypertension. J Heart Lung Transplant 2009;28:373–379.

33. Pousada G, Baloira A, Valverde D. Molecular and clinical analysis of TRPC6 and AGTR1 genes in patients with pulmonary arterial hypertension. Orphanet journal of rare diseases BioMed Central; 2015;10:1.

34. Damico R, Kolb TM, Valera L, Wang L, Housten T, Tedford RJ, Kass DA, Rafaels N, Gao L, Barnes KC, Benza RL, Rand JL, Hamid R, Loyd JE, Robbins IM, Hemnes AR, Chung WK, Austin ED, Drummond MB, Mathai SC, Hassoun PM. Serum endostatin is a genetically determined predictor of survival in pulmonary arterial hypertension. Am J Respir Crit Care Med 2015;191:208–218.

35. Paulin R, Dromparis P, Sutendra G, Gurtu V, Zervopoulos S, Bowers L, Haromy A, Webster L, Provencher S, Bonnet S, Michelakis ED. Sirtuin 3 deficiency is associated with inhibited mitochondrial function and pulmonary arterial hypertension in rodents and humans. Cell Metab 2014;20:827–839.

36. Rhodes CJ, Batai K, Bleda M, Haimel M, Southgate L, Germain M, Pauciulo MW, Hadinnapola C, Aman J, Girerd B, Arora A, Knight J, Hanscombe KB, Karnes JH, Kaakinen M, Gall H, Ulrich A, Harbaum L, Cebola I, Ferrer J, Lutz K, Swietlik EM, Ahmad F, Amouyel P, Archer SL, Argula R, Austin ED, Badesch D, Bakshi S, Barnett C, et al. Genetic determinants of risk in pulmonary arterial hypertension: international genome-wide association studies and meta-analysis. Lancet Respir Med 2019;7:227–238.

