## Supplementary material for "Common variation in *EDN1* regulatory regions highlights the role of PPARγ as a key regulator of Endothelin *in vitro*": Supp. Table

**Supplementary Table 1**: list of primers used for PCR amplification of the whole EDN-1 sequence.

| **Fragment** | | **Primer** | **Sequence** **(5’-> 3’)** | **Tm** **(ºC)** | **PCR product** |
| --- | --- | --- | --- | --- | --- |
| 5’UTR | *EDN-1* P1 | Forward | CATTCCTTGACCCTCCTCGG | 59 | 815 bp |
|  |  | Reverse | CAGGCCCGAAAGGAAATCAC |  |  |
|  | *EDN-1* P2 | Forward | GCAGTGATTTCCTTTCGGGC | 59 | 117 bp |
|  |  | Reverse | GAGTGGGGGTAAACAGCTCC |  |  |
|  | *EDN-1* PE | Forward | TGGGGCTGGAATAAAGTCGG | 58 | 436 bp |
|  |  | Reverse | AAGTCAACGAGCGTGCCTA |  |  |
|  | *EDN-1* EI | Forward | AGAAACAGGTAGGCACGCTC | 56 | 113 bp |
|  |  | Reverse | ACACTTGCTCTCTAAGCTGC |  |  |
| 3’UTR | *EDN-1* F1 | Forward | GTGACCCACAACCGAGCACATTG | 60 | 1506 bp |
|  |  | Reverse | GCATCTCTCACCACACCATCAGC |  |  |
|  | *EDN-1* F2 | Forward | GGGACTTCATCCATTAACTTGGC | 50 | 1723 bp |
|  |  | Reverse | CGTGCTTAACTTAAGTGTTCAGG |  |  |

**Supplementary Table 2:** list of primers used for PCR amplification of the fragments to clone in the luciferase vectors

| **Fragment** | **Primer** | **Sequence** **(5’ -> 3’)** | **Tm** **(ºC)** | **PCR product** |
| --- | --- | --- | --- | --- |
| *EDN-1* 5’UTR | Forward(Nhe I) | AATAAGCTAGCCATTCCTTGACCCTCCTCGG | 58 | 1434 pb |
|  | Reverse (Xho I) | AATAACTCGAGACACTTGCTCTCTAAGCTGC | 58 |  |
| *EDN-1* 3’UTR F2 | Forward (Nhe I) | AAAGCTAGCGGGACTTCATCCATTAACTTGGC | 72 | 1740 pb |
|  | Reverse (Sal I) | AAAGTCGACCGTGCTTAACTTAAGTGTTCAGG | 72 |  |

**Supplementary Table 3:** list of primers used for qPCR amplification of the transcription factors and housekeeping genes.

| **Primer** | **Sequence (5’ -> 3’)** | **Tm (ºC)** | **PCR product** |
| --- | --- | --- | --- |
| *KLF4–F* | GGCACTACCGTAAACACACG | 60 | 140 bp |
| *KLF4–R* | CTGGCAGTGTGGGTCATATC |  |  |
| *PPARγ-F* | GGTGAAACTCTGGGAGATTCT | 60 | 102 bp |
| *PPARγ-R* | CTCTGTGTCAACCATGGTCA |  |  |
| *VDR-F* | TCCTCCTGCTCAGATCACTG | 60 | 107 bp |
| *VDR-R* | AGGGTCACAGAAGGGTCATC |  |  |
| *EDN1-F* | TCTCTGCTGTTTGTGGCTTG | 60 | 103 bp |
| *EDN1-R* | GACTGGGAGTGGGTTTCTCC |  |  |
| *YWHAZ-F* | ATGCAACCAACACATCCTATC | 60 | 178 bp |
| *YWHAZ-R* | GCATTATTAGCGTGCTGTCTT |  |  |
| *ALAS1-F* | AGTGTGAAAACCGATGGAGG | 60 | 140 bp |
| *ALAS1-R* | CGATCATACTGAAAAGTGGAAACAG |  |  |
| *EDN1-porom-F* | CTCGCTGCCTTCTCTCCTG | 60 | 74 |
| *EDN1-prom-R* | AAAGCGATCCTTCAGCCCAA |  |  |
